# Moment-by-moment tracking of naturalistic learning and its underlying hippocampo-cortical interactions

**DOI:** 10.1101/2020.12.09.416438

**Authors:** Sebastian Michelmann, Amy R. Price, Bobbi Aubrey, Werner K. Doyle, Daniel Friedman, Patricia C. Dugan, Orrin Devinsky, Sasha Devore, Adeen Flinker, Uri Hasson, Kenneth A. Norman

**Affiliations:** Department of Psychology, Princeton University, Princeton, NJ, USA; Princeton Neuroscience Institute, Princeton University, Princeton, NJ, USA; School of Medicine, New York University, New York, NY, USA

## Abstract

Every day our memory system achieves a remarkable feat: We form lasting memories of stimuli that were only encountered once. Here we investigate such learning as it naturally occurs during story listening, with the goal of uncovering when and how memories are stored and retrieved during processing of continuous, naturalistic stimuli. In behavioral experiments we confirm that, after a single exposure to a naturalistic story, participants can learn about its structure and are able to recall upcoming words in the story. In patients undergoing electrocorticographic recordings, we then track mnemonic information in high frequency activity (70 – 200*Hz*) as patients listen to a story twice. In auditory processing regions we demonstrate the rapid reinstatement of upcoming information after a single exposure; this neural measure of predictive recall correlates with behavioral measures of event segmentation and learning. Connectivity analyses on the neural data reveal information-flow from cortex to hippocampus at the end of events. On the second time of listening information-flow from hippocampus to cortex precedes moments of successful reinstatement.

---

Humans can learn very quickly. When meaningful content is presented in narrative form, we are able to absorb a substantial amount of information in one shot, that is without repetition or rehearsal^1, 2^. Consequently, when we listen to a story that we have heard before, we can often recall what is about to happen from memory.

This kind of naturalistic learning has recently received more attention in the investigation of our memory system^1, 3, 4^. Naturalistic stimuli engage the brain to a stronger extent than sparse and artificial stimuli^5^ and allow us to study how memory is used “in the wild”, outside of situations where our memory is explicitly being tested. Studying memory in naturalistic settings has also exposed some fundamental theoretical questions that were previously ignored: When participants are given an artificial memory test (e.g., a list of word pairs), it is easy to say when memories should be stored and retrieved (they should be stored during the “study phase” and retrieved during the “test phase”), but the real world is not conveniently divided up into “study phases” and “test phases” – there is just continuous experience, and the field of memory research is just beginning to grapple with the question of when encoding and retrieval take place in the wild. Recent work in cognitive psychology and cognitive neuroscience has started to shed light on these issues. Cognitive psychology has focused on the role of pre-existing knowledge about how situations unfold, so-called schemata^6^, in structuring our continuous experience into discrete events (e.g., a dinner or a phone call). Theories of event segmentation have productively explored how people automatically and spontaneously segment experience into events and how this segmentation impacts how information is remembered later^7–9^. In cognitive neuroscience, work has focused on how hippocampus and neocortex are engaged during processing of naturalistic stimuli. It has long been known that memory for unique experiences (i.e., episodic memory)^10^, relies on the hippocampus as a key structure^11–13^. The hippocampus is thought to be crucial for forming and retrieving associations between memory content, whereas the detailed representation of that content is ascribed to the neocortex^14^. Indeed, the reinstatement of information-rich memories is frequently linked to cortical signatures^15–17^. In the realm of naturalistic paradigms, functional MRI studies have found that event specific patterns from encoding become reactivated in neocortex during retrieval of stimulus material^1, 3, 18^; the hippocampus, on the other hand, becomes more active at the end of events during memory encoding and this hippocampal activity is predictive of subsequent memory retrieval^3, 19, 20^.

These findings suggest an interplay between hippocampus and neocortex under naturalistic conditions that is shaped by the structure of the narrative, and they engender questions about when and how information is exchanged between the two structures. One would expect that information flows from neocortex to hippocampus during learning, potentially timed to the end of naturalistic events^9, 20^. If the hippocampus then initiates the reinstatement of information in the neocortex, one would expect that information-flow from hippocampus to neocortex precedes the reinstatement of mnemonic information. Importantly, the fine-grained temporal nature of these hippocampo-cortical interactions requires the use of a method with high spatial and temporal resolution: These questions can be optimally addressed via electrocorticography (ECoG) that uses concurrent recordings from neocortical and hippocampal sites, while patients are experiencing a naturalistic narrative that contains several event boundaries.

Contrary to experimental intuition, listening to a naturalistic story may provide an ideal handle on episodic memory by leveraging one of its key features, namely prediction about the future. Predictive processing frameworks^21, 22^ suggest that a ubiquitous function of our brain is to predict the future in order to reduce uncertainty in perception. In the context of subsequent exposures to the same sequence of stimuli, the hippocampus has been suggested to drive recall-based prediction of upcoming information in early processing regions^23^. In line with this, the hippocampus has further been implicated in the expectation of upcoming stimuli^24^. An unconstrained listening-paradigm, in which participants listen repeatedly to the same story, can naturally induce such predictive recall for upcoming information from episodic memory.

In ECoG, a well-described correlate of listening to auditory stimuli - and specifically speech - is entrainment of the high gamma frequency band across auditory processing areas^25–27^. Such high-frequency activity has been shown to reflect the mass firing of neural populations at the recording site^28, 29^. High-gamma activity in response to auditory stimuli can also be modulated by top-down information: Melodic expectations, for instance, modify this high-gamma response^30^ and prior experience with a clean version of degraded speech can help to understand the degraded version via rapid tuning of the high-frequency response in the auditory cortex^31^. In other words, early auditory responses can adapt based on contextual information^32, 33^. A hypothesis that follows from these observations is that rapid adjustments of neural responses should also arise from episodic memory after a single exposure to a naturalistic story, thereby allowing for the tracking of cortical memory content in the form of anticipation.

We assessed learning under naturalistic conditions in patients undergoing electrocorticographic (ECoG) recording for clinical purposes and in healthy participants (who did not undergo ECoG recording), as they listened to the short story “Pie Man” by Jim O’Grady. To determine the event structure of the story, we first collected human-annotated event boundaries in healthy participants. Participants then repeated the task which allowed for the assessment of learning based on consensus and response time changes. In a separate set of behavioral experiments, we directly tested the learning of story content. To this end we prompted groups of healthy participants about upcoming words with and without prior exposure to the story^34^. The patients that were undergoing ECoG recording simply listened to the story; after a break, they listened to the story again. We probed for neural signals of predictive recall via Granger Causality analysis by testing whether the amplitude of the high-frequency signal (70 – 200*Hz*) contained more information about upcoming neural states on the second run of listening compared to the first. We then traced the information-flow between cortical electrodes that expressed such predictive recall and hippocampal electrodes, examining the temporal dynamics of hippocampo-cortical interactions in the vicinity of event boundaries. To directly investigate hippocampo-cortical interactions related to the reinstatement of information, we then scrutinized the model predictions from the neural Granger Causality analysis. From this we derived a moment-by-moment time course of predictive recall (i.e., moments when predictive information became available from memory), which we then tested for its relationship to event boundaries and to behavioral prediction learning. Finally, we identified punctuate moments in this time-course of predictive recall, where the neural signal strongly and correctly predicted upcoming moments in the story (i.e., peaks). Hippocampo-cortical interactions in the direct vicinity of these peaks revealed the information-flow between hippocampus and cortex that subserves predictive recall.

## Results

### One-shot learning of human-annotated event boundaries in the story

A crucial advantage of naturalistic paradigms is that they allow for the assessment of continuous structure as it is encountered in real life, enabling us to investigate the role of event boundaries^7^ in learning. To this end, we collected human-annotated event boundaries on Amazon Mechanical Turk^35^. We also used these data to assess whether participants’ perception of event boundaries changes after a single exposure to the story. A total of 205 participants performed a boundary detection task: They listened to the story with the instruction to press the space bar whenever, in their opinion, one natural and meaningful unit ended and another began. After that, they listened to the same story and were asked to do the task a second time. Response data were coded for every participant as a vector that recorded 0 or 1 at every millisecond in the story, indicating whether the space bar was pressed within the surrounding second. The averaged time-course can be interpreted as the degree of agreement on event boundaries, specifically the proportion of participants that perceived each moment in the story as an event boundary (Fig. 1a).

**Figure 1.**
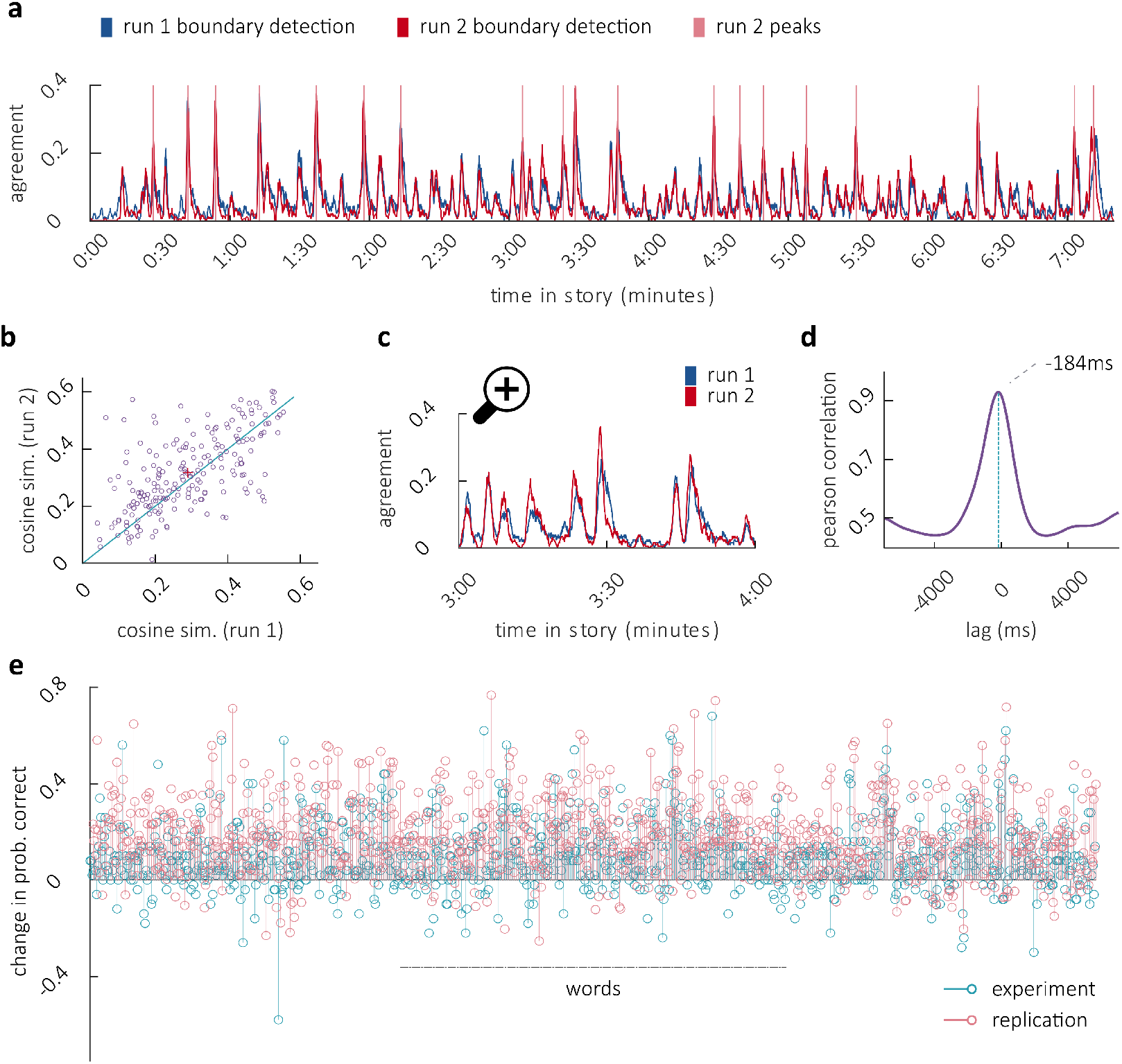
One-shot learning of event boundaries and word predictions. **a.** Agreement between raters on event boundaries. Blue and red lines depict the ratio of raters that marked an event-boundary within every second in the story on run 1 and run 2. Lines in fuchsia indicate boundaries based on run 2, as per peak detection. **b.** Similarity of each subject’s response vector to the agreement between all others. Purple dots indicate individuals: subjects above the diagonal increase in similarity to others on the second run. The average increase (red cross) indicates consensus learning. **c.** Zoomed-in time interval of agreement between raters (compare panel a). Agreement on the second run (red line) slightly precedes agreement on the first run (blue line). **d.** Earlier boundary detection on run 2 is marked by a negative lag in the cross-correlogram (purple), peaking at –184*ms* (turquoise vertical line). **e.** Performance difference between groups that have listened to the story and naive participants in prediction of upcoming words in the story (the naive group was guessing). Predictive recall of upcoming words manifests itself in a positive difference between the groups, i.e. prediction probability of the words increases after a single exposure (turquoise experiment 1, fuchsia replication), demonstrating one-shot learning of story content.

#### Increased consensus upon one-shot learning

We predicted that participants would have a better understanding of the underlying event structure of the story on the second run, and therefore would agree more about event boundaries on the second compared to first run, despite not knowing of others’ responses and not receiving corrective feedback. We analyzed agreement for each participant by computing the cosine-similarity between each participant’s response vector (coded as 0,1) and the average agreement vector between the remaining (*n* – 1) participants (Fig. 1b). Cosine-similarity to others on run 2 (*mean* = 0.318) was significantly higher than on run 1 (*mean* = 0.291, *p* = 0.008 as per 1000 random assignments of run-labels, Supplementary Fig. 1c, right), with 61.46% (126/205) of participants increasing in cosine-similarity (*p* < 0.001, note: increased agreement can also be assessed in the distribution of agreement. This is visible in a quantile-quantile plot and can be assessed via kurtosis, see Supplementary Text and Supplementary Fig. 1a-b)

#### Earlier boundary detection upon one-shot learning

We further predicted that participants would anticipate upcoming event boundaries on the second run and therefore be slightly faster in their responses (compare Fig. 1c). We tested this hypothesis via cross-correlation between the time-course of agreement from the second and from the first run. The maximal cross-correlation (*r* = 0.931) was observed at a negative lag of – 184*ms* (Fig. 1d), indicating that participants became significantly faster in detecting event boundaries (confirmed by 1000 cross-correlations between average time-courses from randomly permuted run-labels, *p* < 0.001, yielding no lag that was more extreme).

### One-shot learning of story content

Next we wanted to directly confirm that one-shot learning behaviorally enables the predictive recall of story content. To this end, behavioral data from 100 previously collected participants that performed prediction experiments were available in aggregated form; we collected an additional 100 participants on Amazon’s Mechanical Turk^35^, to replicate and extend the findings. In both experiments, participants predicted upcoming words in the story from a context of 10 previous words that were presented in written form on their screen (beginning at word 11 in experiment 1 and at word 3 in experiment 2, with less context for the first 8 predictions). Only half of participants (*N* = 50 per experiment) listened to the story before performing the task; the other half had not listened to the story and therefore could not use episodic memory to recall what comes next. After naming the next word, the correct next word was revealed and participants guessed (in the first listening condition) or recalled (in the predictive recall condition) the next upcoming word. Sliding this contextual window along the story generates a prediction-probability score for each word in the story, in each condition. Prediction-probability of the words was higher in the group that had listened to the story in the prediction experiment (*t*(934) = 23.043, *p* < 0.001, *d* = 0.754) and in its replication (**t**(962) = 44.188, *p* < 0.001, *d* = 1.424, Fig. 1e), confirming one-shot learning of content. In the replication, where data were available on the participant level, we could also compare participants’ prediction performance between groups. Participants that had heard the story before predicted more words correctly (mean = 390.24, std = 180.151) than naive participants (mean = 214.6, std = 82.225, *t*(98) = 6.272, *p* < 0.001, *d* = 1.267), confirming again the rapid learning of story content.

### Neural evidence for predictive recall

The behavioral observations of predictive recall on the second run of listening suggest that it should be possible to observe the emergence of predictive information in neural signals during the second presentation of the story. We therefore probed for predictive recall in patients undergoing electrocorticographic (ECoG) recording. By applying Granger Causality (GC) analysis^36–38^ between the first and the second run of listening, we assessed whether there was more information about upcoming states in the amplitude of the high-frequency signal (70 – 200*Hz*) on the second run of listening compared to the first run. GC tests if the past of a signal *Y* (i.e., up to time point *t*, see: Online Methods) can improve the prediction of signal *X* (at time point *t*), above and beyond what the past of signal *X* can predict about its own future states^37^. Typically GC is interpreted as a measure of causal relation; certainly there is no causal influence from the second run onto the first – the logic here is that, if patients are using episodic memory to anticipate what will happen next in the story, information about the story should appear in the neural signal earlier in run 2 than in run 1. Concretely, the past signal of run 2 should predict the future signal of run 1, above and beyond what the past signal of run 1 can predict about its own future states - exactly the circumstance that GC is meant to capture. Crucially, the above hypothesis can be tested by assessing the asymmetry of GC across runs: In the full model, the past of run 2 should predict the future of run 1, more so than the past of run 1 predicts the future of run 2 (Fig. 2a). We further included the envelope of the auditory signal in the full model, which controlled for entrainment from low level stimulus-features and allowed us to test for learning of these features (see below)

**Figure 2.**
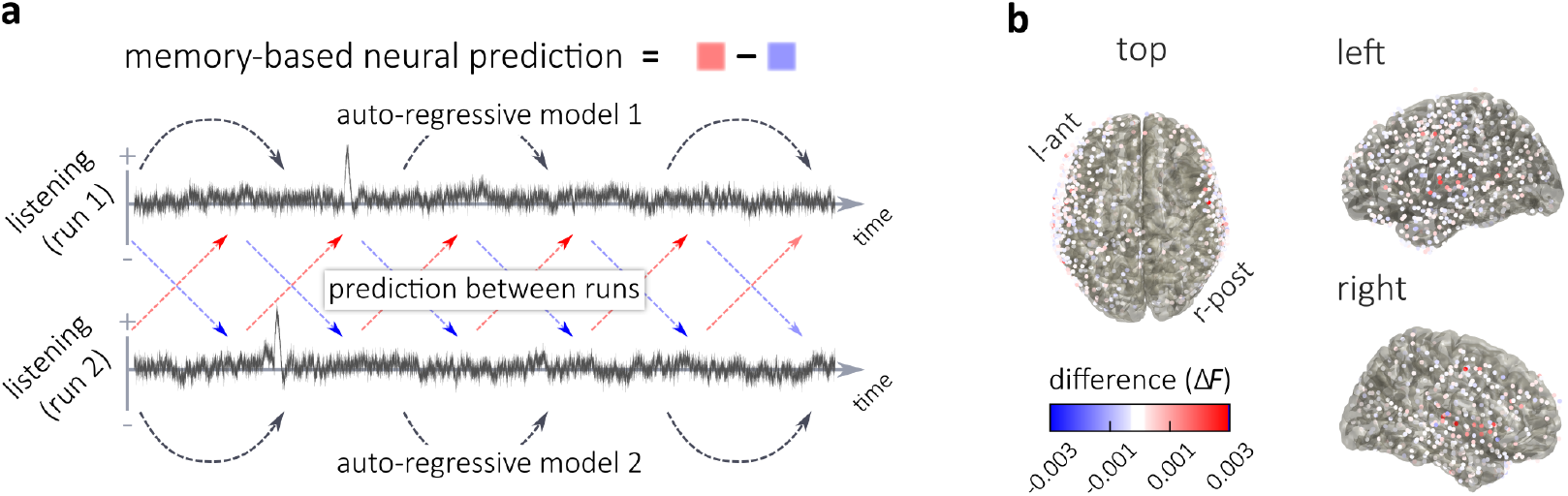
Identifying electrodes that show predictive recall. **a.** Tracking of neural prediction via Granger Causality: Prediction between runs is added to each auto-regressive model. If the neural signal acquires memory about upcoming states after listening to the story once, then data from run 2 should be able to improve the prediction of the auto-regressive model for run 1 (red arrows). Signal from run 1, on the other hand, should not be able to improve the prediction of the auto-regressive model for run 2 (blue arrows). The difference between those predictions across runs is interpreted as a measure of predictive recall. **b.** Difference in *F*-values between the prediction of run 1 from run 2 and the prediction of run 2 from run 1, indicating neural evidence for predictive recall that emerges on the second run of listening

We contrasted how much the second run could improve the prediction of the first run with how much the first run could improve the prediction of the second, by taking the difference in the respective *F*-values (Fig. 2b). This difference, averaged across all electrodes, was significantly smaller when the data were randomly assigned to run 1 and run 2 for each electrode (*p* = 0.001) and for each patient (*p* = 0.005, see Supplementary Fig. 3a). No significant effects were found in other frequency bands (all *ps* > 0.135) and we found no significant evidence for learning of low-level stimulus-information (comparing predictive information about the audio-envelope between runs, *p* > 0.8). To select electrodes on which mnemonic information was present, we fitted a Gaussian mixture model with 2 underlying distributions (one should fit around zero, one not) to the differences in *F*-value across all patients’ electrodes. Thirty-one electrodes, hereafter referred to as ‘cortical predictive recall’ (CPR) channels, were selected for further analyses, because the posterior probability of their observed difference in *F*-value was 10 times higher for the ‘effect distribution’ compared to the ‘null distribution’ (Supplementary Fig. 3b). These electrodes were located in cortical auditory processing regions (Supplementary Fig. 3c); we did not observe predictive recall effects on hippocampal electrodes. Interestingly, the optimal model order for measuring neural predictive recall (taking into account neural activity between up to 130*ms* and 350*ms* as per Akaike Information Criterion, see: Online Methods) was in the time-range of the behavioral advancement of 184*ms* for boundary detection. This invites speculation about an association between predictive information that is available in auditory processing regions and behavioral benefits.

### Hippocampo-cortical interactions near event boundaries

Prior evidence implicates the hippocampus in the processing of events, such that hippocampal activity is increased at the offset of events and the amplitude of such offset responses is predictive of subsequent recall performance^3, 19^. Based on these results, we predicted information-flow from CPR-channels to the hippocampus at the end of events. We tested this hypothesis in patients where hippocampal channels were recorded and predictive recall was observed (*N* = 6): For every hippocampal channel, multivariate mutual information (MI) with CPR-channels was assessed, which measures statistical dependence between the channels and quantifies shared information. Specifically, we concatenated the high-frequency amplitude (70 – 200*Hz*, Fig. 3a top) on the CPR-channels within 1 second around the 19 potential event boundaries (identified using the button-responses in the behavioral experiment’s run 2). For each hippocampal channel, we concatenated the amplitude across 6 frequency bands (delta < 4 Hz, theta 4 – 8, alpha 8 – 15, beta 15 – 30, low gamma 35 – 55 and high gamma 70 – 200) and the raw signal (Fig. 3a bottom), in order to be sensitive to any shared information. To account for delay in information-flow between channels, we assessed MI repeatedly at different lags (between a 1.5*s* lead and 1.5*s* lag), by shifting the channels in steps of 10*ms*. Furthermore, we conditioned MI on the (typically spurious) shared information at zero lag (signal not shifted)^39, 40^. Finally, because there is a perception-to-action delay between the moments in which patients perceive an event boundary and the moment in which they indicate it using the button press, we repeated this whole analysis at all potential moments around the marked time of event boundaries (–3*s* to +1*s*). This analysis therefore resulted in a map of shared information between hippocampus and the CPR-channels at each channel-lag (conditioned on the zero lag) and at varying distances from the event boundary.

**Figure 3.**
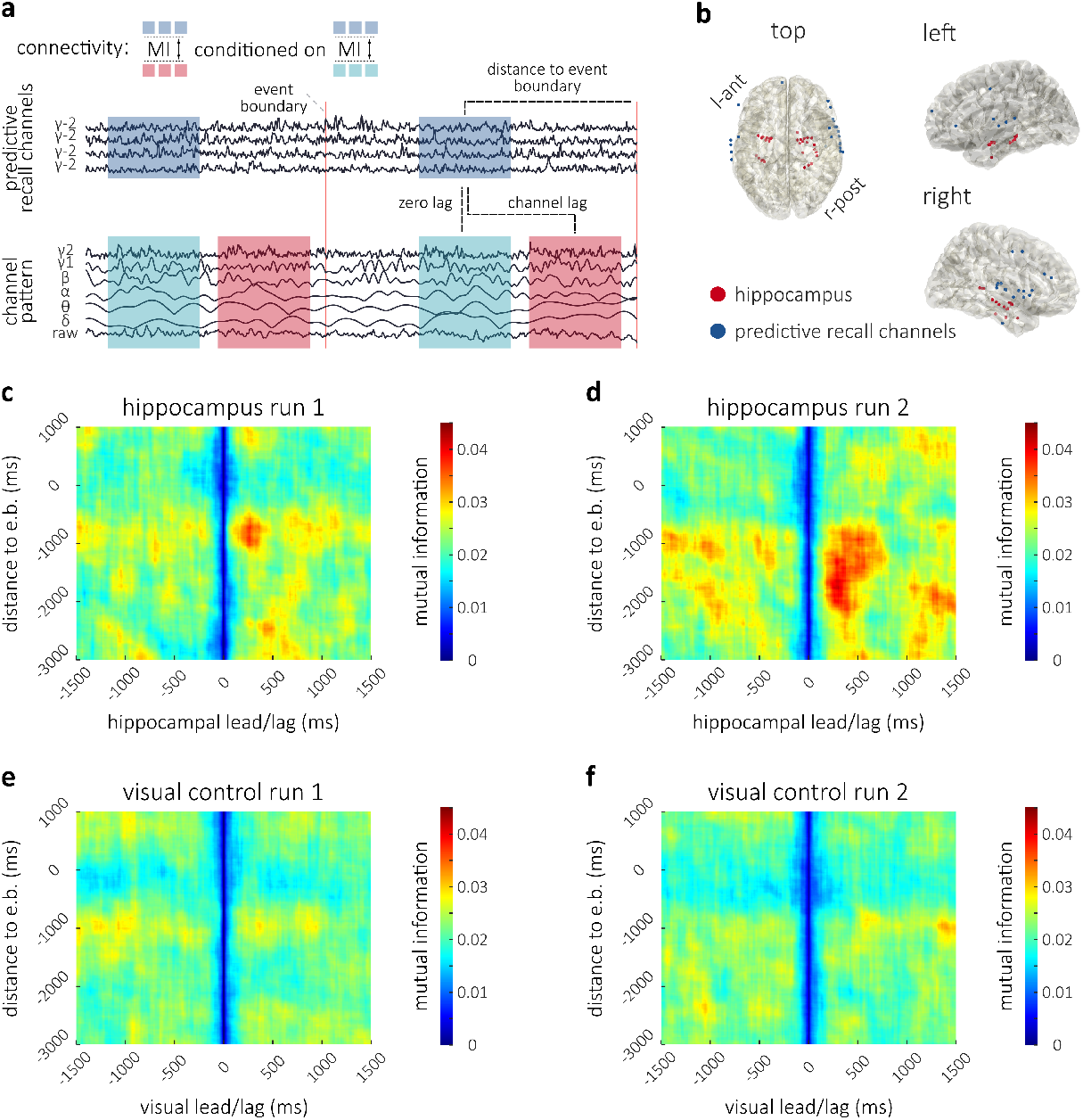
Connectivity analysis between ‘CPR-channels’ and hippocampus/visual control. **a.** Conditional Multivariate Mutual Information was computed across 1s time-windows between the high-gamma amplitude from all available ‘CPR-channels’ (blue) and 6 frequency bands and raw data (red) at each respective channel of interest. This was done at different channel-lags; the analysis at each lag was conditioned on the zero-lag pattern (turquoise). This analysis, which yields an estimate of shared information at each lag, was repeated at different distances to event boundaries (red lines), resulting in a 2 dimensional map. **b.** ‘CPR-channels’ (blue) and hippocampal electrodes (red). **c.** Run 1 map of MI between hippocampus and ‘CPR-channels’ at different lags and distances to boundaries. MI peaks at 730*ms* before event boundaries (button press, y-axis); at the peak, information in hippocampus lags behind ‘CPR-channels’ by 270*ms* (x-axis). **d.** Run 2 map displaying an earlier peak at 1770*ms* before boundaries with a 300*ms* hippocampal lag. **e.** Run 1 map of MI between electrodes in a visual control ROI and ‘CPR-channels’ at different lags and distances to boundaries. **f.** Run 2 map of MI between electrodes in a visual control ROI and ‘CPR-channels’ at different lags and distances to boundaries.

At a distance of 730*ms* before event boundaries (button-responses in the behavioral sample, note that patients were only listening), we observed increased MI between hippocampus and CPR-channels at a hippocampal lag of 270*ms* (peak in map), when patients listened to the story for the first time (Fig. 3c), indicating information-flow from CPR-channels to the hippocampus (compare Fig. 3b). To assess significance of this information-flow, we compared this connectivity profile to the map of MI between the same CPR-channels and a control region of interest in visual cortex (network 1 from Yeo atlas^41^: 31 channels, see Supplementary Fig. 2b) where we did not expect information-flow (Fig. 3e). For multiple comparison correction across multiple lags and time points, we used a cluster-based permutation approach (cluster-forming threshold 95*^th^* percentile in 2-sample test between regions, see Online Methods). In this analysis, we considered clusters of neighboring time/lag-points within a plausible time period around the boundaries (2*s* prior to 500*ms* post button-response) and a plausible range of lags (1*s* lead to 1*s* lag); note that a wider range was computed and plotted for examination at the reader’s discretion. The summed maximum cluster in the hippocampal analysis (peaking at 730*ms* before the button-response at 270*ms* hippocampal lag) was significantly greater (*p* = 0. 044) than maximum cluster-sums under 1000 random swaps of channels between the regions on the first run of listening.

We next investigated MI between hippocampus and CPR-channels when patients were listening to the story for the second time. Again we found enhanced information-flow from CPR-channels to hippocampus near event boundaries (peaking at 1770*ms* before the button-response at a hippocampal lag of approximately 300*ms*, Fig. 3d). Statistical comparisons confirmed that the maximum cluster was again larger for the hippocampal channels than for the visual control ROI (*p* = 0.017, Fig. 3f). Interestingly, on the second run of listening, there was more MI between hippocampus and CPR-channels, and information-flow was observed earlier than on the first run of listening (peaking at 1770*ms* compared to 730*ms* before the button-response). A comprehensive summary of these findings is that the ends of events are important moments for memory encoding: when patients anticipate an event boundary, enhanced information-flow from cortex is initiated earlier and more robustly.

### Moment-by-moment tracking of predictive recall

The above findings describe information-flow from cortex (CPR-channels) to hippocampus in the vicinity of event boundaries. Event boundaries, however, represent moments of high uncertainty, where information about upcoming states is sparse^9^. One would therefore expect predictive recall to minimize this uncertainty via information-flow from hippocampus to cortex, when information is available in episodic memory. While we did not directly observe significant evidence for such information-flow from hippocampus to cortex during the second exposure in the vicinity of event boundaries, the use of human-annotated event boundaries may arguably not provide the best handle to predictive recall itself: Even if predictive recall were taking place around event boundaries, the boundaries are an aggregate measure (derived from a separate group of participants) and thus may not capture the potentially-variable timing of predictive recall in individual patients. We therefore directly asked the question: which moments in the story contributed to predictive recall in individual patients? To answer this question we interrogated the model predictions between runs from the Granger Causality analyses (compare Fig. 2) within each patient: A predicted signal of run 1 was derived from run 2, using the coefficients from the GC-model. These predictions were then projected onto the actual data at run 1. As a contrast, run 2 was predicted from run 1 and projected onto the data from run 2. The difference between these projected model-predictions represents a moment-by-moment measure of predictive recall. Peaks in this time-course can be interpreted as moments that become substantially more anticipated by patients on the second run of listening, in other words, moments of strong and correct predictive recall. These points correspond to meaningful moments in the story, for instance profanity and humor (Fig. 4a, see: Supplementary Information for a video).

**Figure 4.**
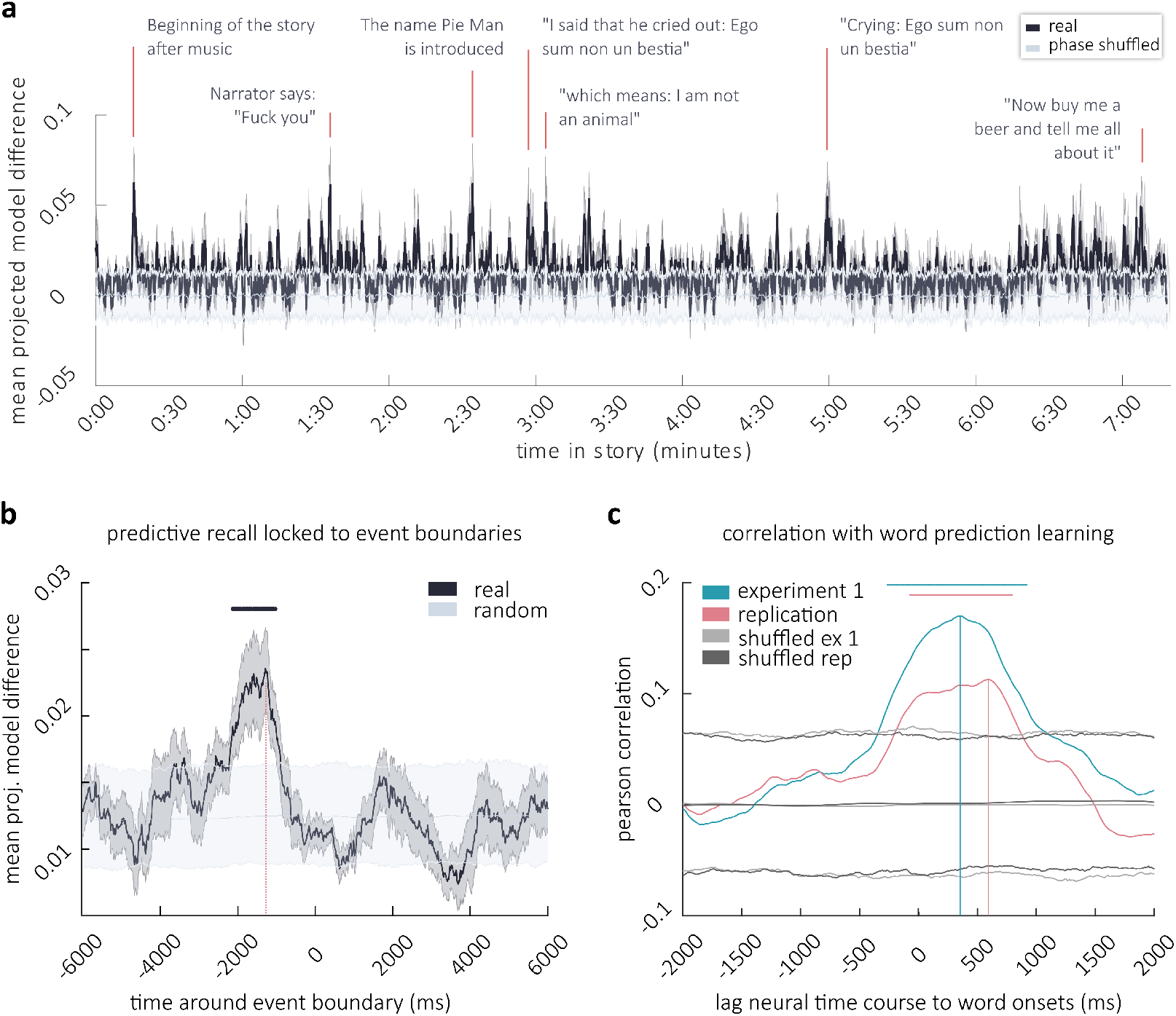
Moment by moment tracking of predictive recall throughout the story and relation to behavior. **a.** Difference in model predictions projected onto the data at every moment in the story (Black, ±SEM gray). At peaks the model from run 2 matches the data on run 1 substantially better than vice versa. Applying the coefficients to phase-shuffled data renders the neural prediction meaningless (light gray, error-bars are 5*^th^* and 95*^th^* percentile). **b.** Neural time-course of predictive recall (a) locked to event boundaries from run 2 (black line ±*SEM*). The gray line depicts mean, 5*^th^* and 95*^th^* percentile of neural data averaged 1000 times across random boundaries. Horizontal black line marks significance, the vertical line in fuchsia marks the peak. **c.** Correlation between the behavioral measures of increase in word-prediction performance through learning (compare Fig. 1e) and the neural time-course of predictive recall (a) at different time-lags. Lines in turquoise and fuchsia show correlation with data from behavioral experiment 1 and replication, respectively. Gray lines are 5*^th^* and 95*^th^* percentile of correlations under random assignment of the change in prediction probability to individual words. Horizontal lines mark significance, vertical lines mark peaks.

### Neural predictive recall relates to event boundaries

Behavioral learning about event boundaries suggests that neural predictive recall will encompass the structure of the story and allow, for instance, the anticipation of boundaries. We tested this hypothesis by analyzing the time-course of memory-based neural prediction directly around 19 event boundaries (local peaks, extracted from the time-course of agreement on the second behavioral run). We averaged 19 segments of neural prediction data around these boundaries and compared it to averages derived from 1000 random selections of those 19 boundaries (Fig. 4b). The increase in memory-based prediction exceeded multiple comparison corrected chance level at several time points between 2140*ms* and 1020ms before event boundaries (i.e., button-responses), with a peak at –1290*ms* (see Supplementary Information for a fine grained analysis via cross correlation); note that this negative lag is expected because a behavioral response can only happen after an event has been neurally registered. These data provide evidence that neural prediction learning encompasses anticipation near event boundaries (i.e., right before button-responses were given in the behavioral sample).

### Neural predictive recall reflects one-shot learning of story content

We further expected that the neural time-course of predictive recall would reflect what is being learned about individual words. To assess this, we correlated the strength of behavioral prediction learning (i.e., the word-level change in prediction-probability, Fig. 1e) with the neural time-course of predictive recall (i.e., the mean projected model difference, Fig. 4a) at all those moments where a word was presented (signal between word-onset and word-offset, excluding silences). We accounted for the latency between word-presentation and neural activity by repeating this analysis under different shifts of the time axis (from 2 seconds before to 2 seconds after word onset). This resulted in a correlation coefficient at different times around the word onset. To assess significance, we conducted the same analysis but assigned the difference in prediction-probability (i.e. behavioral predictive recall) randomly to the words (1000 permutations). After correction for multiple comparisons, the difference in prediction-probability was significantly correlated with the neural data at multiple lags in the first experiment (*p_FDR_* = 0.015, controlling the false positive rate at (*q* = 0.05) with all significant p-values smaller than *p_FDR_*) with a maximum correlation of *r* = 0.17 at 350*ms* after word onset. In the replication (after exclusion of outliers: 9 naive, 7 learning, whose correct responses - coded as 0 and 1 - had a cosine-similarity < 0.6 to the average response accuracy across all other patients), the maximum correlation between neural and behavioral predictive recall (*r* = 0.113) was found at 590*ms* after word onset (*p_FDR_* = 0.006, Fig. 4d, see Supplementary Information for a more sensitive analysis yielding a peak at 320*ms*). These peaks in cross-correlation are in the vicinity of the peak neural high frequency response (70 – 200Hz) to word-onset (peaking at 320*ms*, run 1 and at 389*ms*, run2, see Supplementary Information), suggesting that neural learning entailed the anticipation of individual words by forecasting the high frequency response to word-onset.

### Hippocampo-cortical interactions near predictive recall events

Having identified a moment-by-moment measure of predictive recall, we next investigated the shared information between hippocampus and cortex directly at punctuate moments of high predictive recall in patient-specific time-courses: We first derived an average time-course for every patient and then identified local peaks. Because the exact onset of these neural predictive recall events could be accurately estimated from the data (without relying on a separate set of annotators), we did not need to repeat this analysis at different distances to the peak. This analysis therefore results in a 1 dimensional estimate of shared information at different channel-lags (shifted again in steps of 10*ms*). On the first run, we expected information-flow from CPR-channels to hippocampal channels after predictive recall events (i.e., reflecting the encoding of information). On the second run (but not the first run), we hypothesized that information-flow from hippocampus to the CPR-channels would precede peaks in predictive recall. A chance distribution was obtained by phase-shuffling the neural predictive recall time-courses 1000 times before selecting peaks; p-values (– 1500*ms* lead to 1500*ms* lag) were corrected for multiple comparisons at different lags with a false discovery rate correction^42^. Additionally, the MI between CPR-channels and the hippocampus was compared to the MI between CPR-channels and other channels (all channels that were not CPR-channels or hippocampal channels): We predicted that hippocampus in particular would show the pattern of enhanced information-flow to the CPR-channels in cortex.

In line with our hypothesis, on the first run, no significant information-flow from hippocampus to cortex was observed (Fig. 5a, left). Neither, however, did we find information-flow from cortex to hippocampus in run 1 at these moments in the story that (during run 2) yielded high predictive recall. This suggests that, under naturalistic conditions, information may not always be encoded immediately after it is encountered. Rather, the ends of events may be crucial time-windows for the encoding of information in continuous narratives (compare Fig. 3). On the second run, we did find information-flow from hippocampus to CPR-channels at moments of predictive recall. At lags between –750*ms* and –710*ms* and between –710*ms* and –690*ms* (peak at –740*ms*, negative lags indicate hippocampus precedes cortex) mutual information between hippocampus and CPR-channels was significantly higher than in phase-shuffled data (*p_FDR_* = 0.006) and significantly exceeded mutual information between other channels and the CPR-channels (*p_FDR_* = 0.005, Fig. 5, middle). MI at these lags was also higher on run 2 compared to run 1 (*p_FDR_* = 0.011, as per a dependent sample t-test between runs) and the difference in the real data was significantly higher than differences in phase-shuffled data (*p_FDR_* = 0.004, Fig. 5b). We explored whether any other region (defined as a network in the Yeo atlas^41^, compare: Supplementary Fig. 2b) expressed enhanced information-flow to the CPR-channels, however, we found no significant effect in any other network on the second run of listening. This uniquely links the hippocampus to predictive recall: when predictive recall takes place information flows from hippocampus to cortex.

**Figure 5.**
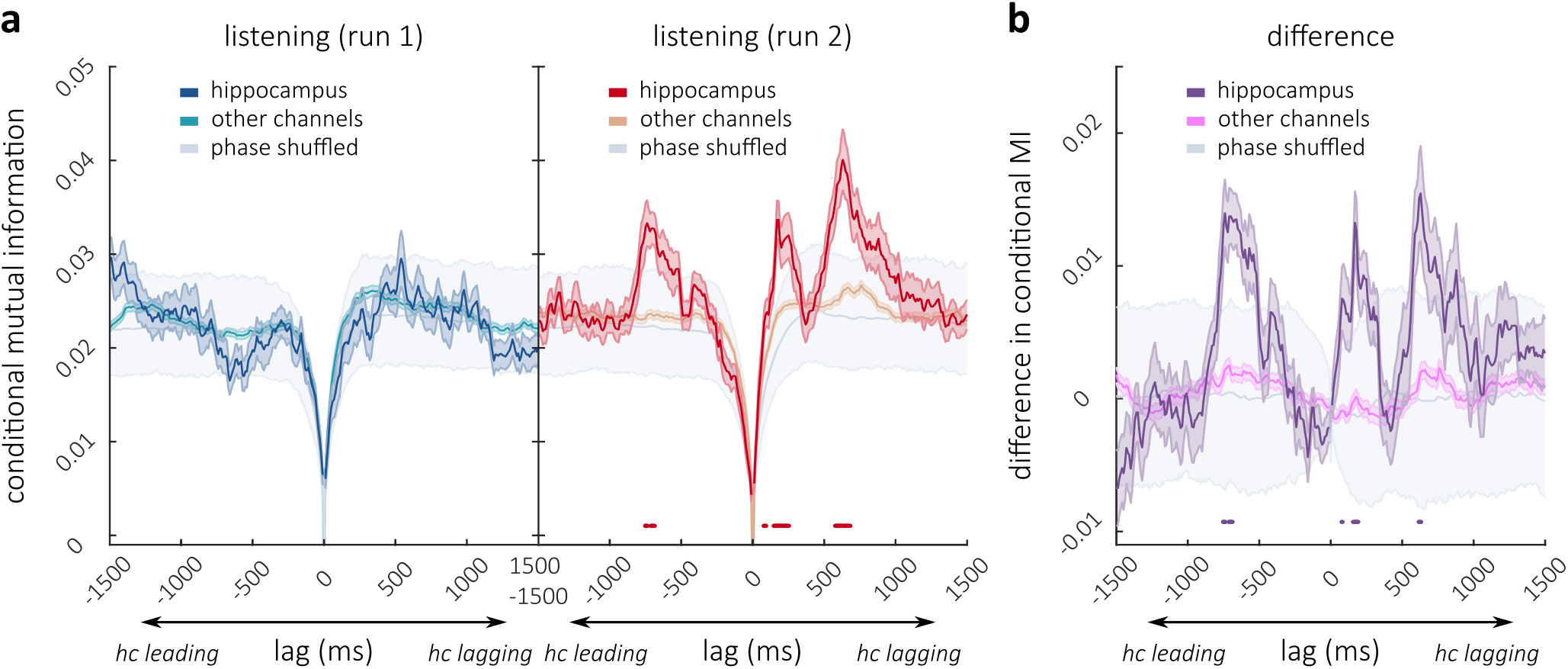
Connectivity analysis at moments of predictive recall. **a.** Conditional Multivariate Mutual Information at different lead/lag to Cortical Predictive Recall channels at peaks in predictive recall on run 1 (left, blue) and run 2 (right, red). MI is conditioned on zero-lag connectivity: a lead signifies information-flow from hippocampus to cortex; a lag signifies the reverse. On run 2 connectivity with hippocampus was significantly increased at prediction events. This was the case at a hippocampal lead of approximately 700*ms* and at 2 different lags. Horizontal red lines depict points of conjunct significance comparing hippocampus to other channels and also to a version of this analysis that uses peaks from phase shuffled neural-data (gray). **b.** Difference in MI between the runs (hippocampus: purple, other channels: pink, phase-shuffled; gray). Horizontal purple lines (right) mark additional conjunct significance of higher connectivity on run 2 compared to run 1, higher difference with real than with phase-shuffled data, and higher difference on hippocampal channels than on other channels (separately FDR-corrected per contrast).

Interestingly, at moments of predictive recall we also found significant evidence for information-flow from ‘CPR-channels’ to hippocampal channels on the second run of listening; this occurred at two distinct time windows: at lags between 80 – 90*ms*, 150 – 250*ms* (peak at a 170*ms*) and again between 580 – 680*ms* (peak at a 630*ms*) information-flow between CPR-channels and hippocampus significantly exceeded shuffled data and the mutual information between other electrodes and CPR-channels (Fig. 5a, right). At lags of 80*ms*, 160 – 190*ms* and 620 – 630*ms* this information-flow was additionally higher on run 2 compared to run 1 (*p_FDR_* = 0.011) and this difference was higher than differences in phase-shuffled data (*p_FDR_* = 0.004, Fig. 5b, see Supplementary Information for a feature-by-feature dissection of these effects).

Taken together, these data shed light onto the fine-grained mechanisms that sub-serve the predictive recall of a naturalistic story. Information from the hippocampus is transferred to cortex at moments of prediction, and information also flows from cortex to hippocampus at those moments, possibly reflecting either feedback signals about successful reinstatement or the re-encoding of information.

## Discussion

These findings unveil the behavioral and neural dynamics at work when we learn under naturalistic conditions. In line with previous experiments, we found that participants perceive boundaries between discrete events when they listen to a story, and we show that these boundaries become anticipated after a single experience. Using connectivity analysis on neural data from patients undergoing ECoG recording, we observed information-flow from auditory cortex to hippocampus at the end of events. This provides novel ECoG support for the idea, derived from fMRI work by Baldassano et al.^3^ and Ben-Yakov and Dudai^19^, that the hippocampus encodes ‘snapshots’ of cortical activity at event boundaries. When investigating predictive recall directly, we found evidence in both behavioral and neural data that predictive information emerges after one-shot learning of the story. We quantified neural predictive recall via Granger Causality analysis, leveraging the fact that recall of upcoming information (during run 2) makes it possible to use neural data from run 2 to predict the future of run 1, but not vice-versa. While there was no explicit instruction for the intracranial patients to memorize the story, neural predictive recall was still reflective of our behavioral measures of learning. Our data therefore constitute a unique demonstration of learning as it naturally occurs without the motivated memorization that typically takes place in memory experiments. Crucially, we identified specific moments in the neural time-course of predictive recall in individual patients when strong anticipatory information emerged from memory. When we time-locked our analysis to these specific moments, we were able to show that information flows from hippocampus to auditory cortex when predictive recall takes place, followed by information flow from cortex to hippocampus.

The moment-by-moment tracking of memory in ECoG patients demonstrates the rapid neural changes that occur with one-shot learning: high frequency activity in auditory processing regions acquires predictive information about upcoming states. Previous work has already shown that high frequency activity in the auditory cortex can tune to auditory input, enhancing the intelligibility of distorted speech through experience^31^. Furthermore, content-specific patterns of activity in gamma power have been linked to the reinstatement of information during successful spatial navigation^43^ and the viewing of images^44^. Our study leverages this phenomenon and demonstrates (on a moment-by-moment timescale) when information in the auditory cortex naturally reappears in a way that is predictive of upcoming states. Notably, we did not observe these prediction effects on hippocampal electrodes themselves – i.e., the hippocampus did not significantly predict its own future states. However, we did implicate the hippocampus in evoking predictive information about cortical states. This is in line with the idea that a hippocampal index reactivates information in cortical areas^45^.

In keeping with prior work, our data suggest that the boundaries between events are important anchor-points for one-shot learning. Conceptually, event boundaries represent moments of high uncertainty, where information about upcoming states is sparse^9^. The end of an event may therefore be an ideal moment to store a coherent picture before an imminent change in the environment. In line with previous data from a non-naturalistic word memorization task showing that high-frequency activity in the neocortex couples to hippocampus during encoding^46^, we observe such coupling during naturalistic learning in the vicinity of event boundaries^3, 19, 20^. Additionally, when patients were listening to the story for the first time, we did not observe information-flow from cortex to hippocampus directly after moments that subsequently expressed high predictive recall; instead, this information-flow was only observed near event boundaries, further suggesting that event boundaries are crucial moments for memory encoding.

Strongly encoding transitions between events may also enable our memory system to bridge uncertainty^47^: The behavioral consensus learning (i.e. the one-shot increase in agreement on event boundaries) and the anticipation of event boundaries in the story, as well as the correlation between neural prediction learning and agreement on event boundaries, all support the notion that event boundaries are important moments in the story. A prior study using scalp-EEG recordings in a non-naturalistic setting also found reactivation of patterns from the previous event at boundaries in streams of static images^48^. A possible explanation of these findings is that information from an event is rapidly recapitulated at event boundaries when thorough encoding takes place and a snapshot is stored in the hippocampus.

When patients listened to the naturalistic story for the second time, we found information-flow from hippocampus to cortex right before moments of high predictive recall. These data are in line with prior evidence suggesting that, during spatial navigation, the maintenance of object information (here cuing for a goal location) is coupled to hippocampus^49^. While we could link predictive recall indirectly to event boundaries by showing a systematic relationship between the two (compare: Fig. 4b), we did not directly observe significantly enhanced information-flow from hippocampus to cortex near event boundaries (compare: Fig. 3c-d). As noted earlier, this null result may simply reflect the fact that human-annotated event boundaries are an aggregate measure (derived using a separate group of participants), and thus may not be sensitive to variance in the timing of individual patients’ perception of boundaries. Furthermore, the annotations reflect the time of participants’ behavioral response to the boundary but not the moment that the boundary was neurally detected - taken together, these factors mean that the event boundary annotations may not be temporally precise enough to identify brief bouts of communication between hippocampus and cortex. By contrast, our neural measure of predictive recall (based on Granger Causality) can be computed based on each individual patient and thus provides a more precise temporal ‘handle’ to moments where episodic memory enables prediction. This may help to explain why we were able to identify hippocampo-cortical communication when time-locking to these moments.

Nevertheless, these data engender questions about the involvement of event boundaries in the retrieval process. A limitation of our investigation of event boundaries in the prediction process is that patients did not have to think far ahead in the story, possibly limiting reinstatement to immediate information. The role of event boundaries in the reinstatement of information is potentially better addressed in a task that probes the active reinstatement of continuous information over longer periods of time. Evidence from the reinstatement of continuous stimuli already suggests that event boundaries may structure retrieval, serving as stepping-stones through longer memories^50^. It is therefore possible that information is encoded near the boundaries to the next event and that these anchor-points are used in the retrieval process to reconstruct the unfolding of continuous experience.

Overall, these results provide a moment-by-moment window into the emergence of memory after a single exposure to a story, showing the intricate dialogues between hippocampus and neocortex that allow us to use memory to anticipate how a story will unfold, and how these dialogues are shaped by the structure of the story. Our results demonstrate the importance of event boundaries in encoding new information from neocortex into hippocampus; our methods also allowed us to identify specific moments of predictive recall in individual subjects, and we showed that these moments are associated with information transfer from hippocampus into cortex. Taken together, these methods and findings provide new insight into how episodic memory encoding and retrieval work in naturalistic conditions at a fine temporal scale.

## End Notes

## Acknowledgements

This work was supported by R01 MH112357 awarded to Uri Hasson and Kenneth A. Norman. The authors wish to thank Janice Chen, Arianna Zuanazzi, Qihong Lu and Ariel Goldstein for helpful discussions, B. Mahmood, L. Fanda, M. Hofstadter, I. Davidesco, A. Zadbood, A. Rao, and P. Minhas for help with with data collection, H. Wang for electrode reconstruction, and Elizabeth McDevitt and Chi Thao “Zoe” Ngo for comments on an earlier version of this manuscript.

## Author Contributions

Amy R. Price and Uri Hasson designed the ECoG experiment and the word prediction task. Amy R. Price collected the online prediction experiment 1 and the ECoG recordings. Bobbi Aubrey collected the replication of the behavioral prediction experiment. Sebastian Michelmann and Kenneth Norman designed the behavioral segmentation experiment. Sebastian Michelmann, Uri Hasson and Kenneth A. Norman analyzed and discussed the data and wrote the manuscript.

## Online Methods

### Stimulus material and experimental procedures

#### Stimulus material

The stimulus material consisted of a humorous story of 7 minutes and 30 seconds duration (“Pieman” by Jim O’Grady), recorded at a live performance (“The Moth” storytelling event, New York City). For the online prediction experiments, the story was transcribed manually by 2 different transcribers for experiment 1 and the replication; a difference in word count between the transcripts is due to the use of hyphenated words (e.g., working-to-middle-class) in experiment 1. Laughter, breathing, lip-smacking, applause, and silent periods were also marked in order to improve the accuracy of subsequent alignment. To derive accurate word onset and offset times, the audio was downsampled to 11 kHz and aligned to the transcription via Penn Phonetics Lab Forced Aligner^51^.

#### Online experiments

A total of 405 healthy volunteers participated in online experiments on Amazon’s Mechanical Turk^35^. Participants provided informed consent before participation in accordance with the Princeton University Institutional Review Board. 205 participants completed a task that required the segmentation of the story into natural and meaningful units; 200 participants completed a different task that required the prediction of words in the story in 2 separate experiments (100 per experiment; the second experiment replicated the first one). The segmentation experiment asked participants to segment the story while they were listening to the audio recording. The verbatim instruction was: “press the space bar every time when, in your judgment, one natural and meaningful unit ends, and another begins”^7^. During the task, a black dot appeared on the screen whenever the space bar was pressed. After completing this task, participants were informed that they would now hear the same story a second time and were asked to segment the story again.

The prediction experiment had been previously run as part of a separate study and data were available in aggregated form; it probed how well participants could predict each word in the story without having previously heard the story, and how well participants could predict each word after listening to the story once^34^. A second experiment replicated the first prediction experiment and collected information about individual word predictions that was no longer available from Experiment 1. In both prediction experiments, participants saw 10 words from the transcribed story presented on their screen. They were asked to predict the next word that followed; after typing their response on their keyboard, the correct next word was shown and participants were asked again which word followed the current set of 10 words (continuing for every word in the story). In experiment 1, this task started at word 11, in the replication the task started at word 3, limiting the initial context. Importantly, in each experiment, 50 participants completed the prediction task without knowing the story, and a separate group of 50 participants completed the same task after listening to the audio recording of the story once. This resulted in a total of 50 participants per experiment that guessed the upcoming word without memory for the story, and a total of 50 participants per experiment that could recall the upcoming word from episodic memory.

#### Electrocorticography experiments

Nine patients (18-58 years old, 4 female, 5 right handed, 1 unknown handedness, see Supplementary Table for further information) were recorded at the Comprehensive Epilepsy Center of the New York University School of Medicine. Patients had been diagnosed with medically refractory epilepsy and were undergoing intracranial recording for purely medical purposes. They provided informed consent in oral and written form before participation, in accordance with the Institutional Review Board at the New York University Langone Medical Center. Patients were informed that participation was unrelated to their medical treatment and that they could withdraw their consent at any point without affecting their care. For patients, the experiment consisted of listening to the story twice. Seven patients completed the first and second run of listening on the same day, with a short break ranging from 1 minute to 1 hour and 40 minutes. One patient completed the second run 2 days later. Patient 8 was recorded on the day of implantation once and completed 2 further runs on day 5 after implantation. Because of a different trigger setup, run 1 and 2 could not be aligned for patient 8. Consequently, run 2 and 3 from day 5 were used in lieu of run 1 and run 2. Importantly, this patient did not have any hippocampal recordings and was not included in the connectivity analyses. Furthermore, none of our findings hinge on the inclusion of this patient.

### Behavioral data analysis

#### Behavioral analysis of word predictions

Prediction probability was derived in the behavioral prediction experiment and in its replication as the proportion of participants that predicted the upcoming word correctly. Individual word predictions were considered as correct if the participant’s lower-case text input matched the correct next word in the story. Typos and spelling mistakes were corrected before prediction accuracy was computed.

#### Behavioral analysis of story segmentation

Data from the story segmentation task were aggregated in response vectors at a resolution of 1000 *Hz*. The response vectors were set to 1 if a given participant had pressed the space bar within 1 second surrounding the time point and were set to 0 otherwise. To derive the time-courses of agreement, these response vectors were averaged across participants. In order to test whether the consensus between participants increased, a measure of consensus was computed by assessing the cosine similarity between a participant’s response vector and the average response vector across all other participants. Therein, the average similarity to other’s response and the number of participants that increased in similarity were assessed. In order to test whether responses on the second run happened earlier than on the first run, the cross-correlation between the average response vector from run 2 and from run 1 was assessed and we noted the lag that maximized the correlation.

### Neural data processing

#### Neural data acquisition and electrode localization

Data were recorded from grid arrays (8 × 8 contacts, 10 or 5 mm spacing), linear strips (1 × 8/12 contacts), or depth electrodes (1 × 8/12 contacts) using a NicoletOne C64 clinical amplifier (Natus Neurological, Middleton, WI) with an online reference to a two-contact subdural strip near the craniotomy site. Data were filtered with an analog band-pass filter (pass-band: 0.16-250 Hz) and digitized at a sampling frequency of 512 Hz. Electrophysiological data were available and could be localized for a total of of 1032 channels (95 – 124 channels per patient, see: Supplementary Fig. 2a). 44 of those channels were excluded from analysis due to poor signal quality (0 – 15 channels per patient, see Supplementary Tables). Localization of electrodes was done via the method described by Yang and colleagues^52^ using the co-registered pre-surgical and post-surgical T1-weighted MRIs of each patient. Finally, non-linear transformations of MRIs onto the Montreal Neurological Institute (MNI) MNI-152 template were computed to derive electrode locations in (MNI) space. Electrodes were labelled anatomically based on the 17-network solution by Yeo and colleagues^41^: An anatomical label was derived for each electrode based on the Euclidean distance to the nearest network in MNI-space (see: Supplementary Fig. 2b). For analyses of the hippocampal region of interest, electrodes from 6 patients were identified via manual inspection in MRIcroGL^53^. The electrodes were selected with a liberal criterion, namely if their surrounding signal drop included at least part of the hippocampal structure; thereby 26 depth electrodes and 4 strip electrodes that were lying on the medial side of the temporal lobe were selected (2-8 electrodes per patient, compare: Fig. 3b, red electrodes and Supplementary Tables).

#### Preprocessing of neural data

Electrophysiological data were analyzed in MATLAB 2019a (MathWorks) using the FieldTrip toolbox^54^. Data were cut from 5 seconds before the beginning of the story to 5 seconds after its end. In a first step, channels were rejected based on visual inspection (0 – 15 channels per patient) and moments where artifacts were present, were marked manually. Subsequently, Independent Component Analysis (ICA) was used to reject reference noise^55^. To this end, ICA filters were computed on artifact free data. For ICA computation, data were additionally filtered with a band-stop filter (stopband: 55 – 65, 115 – 125, 175 – 185). The recording was then interpolated within 150ms around moments where artifacts had been marked manually and the amplitude of a channel exceeded 3.5 interquartile ranges above its median. The computed ICA solution was then applied to the interpolated data and between 1 and 3 of the spatially broadest components were rejected manually. Finally, line noise was filtered out (stop-band: 55 – 65, 115 – 125, 175 – 185). All filters were realized with a zero-phase lag 4th order Butterworth IIR filter as implemented in the fieldtrip toolbox^54^, interpolation was done via Monotone Piecewise Cubic Interpolation^56^.

#### Granger Causality analysis

Pairwise-conditional time-domain Granger causality (GC) analysis was realized with the Multivariate Granger Causality toolbox^37, 38^. GC was computed at each channel separately between run 1 of story listening and run 2 of story listening, conditioned on the amplitude of the audio recording (note that this is different from the traditional use of GC in neuroscience that assesses connectivity between channels). To derive the neural signal, the data were band-pass filtered in the frequency range of interest and the amplitude was computed as the absolute value of the Hilbert transform. The audio recording was bandpass filtered between 200 and 5000 Hz and its amplitude was likewise computed via the absolute value of the Hilbert transform. Subsequently, all time-courses were downsampled to 100Hz. The neural data were truncated to the duration of the auditory recording and the linear trend was removed. Finally, in order to reduce the influence of outliers, those moments where the neural data exceeded 5 interquartile ranges above the median were interpolated with a padding of 5 sampling points around the peak; note that this is mostly relevant for the high frequency band (70-200 Hz) where residual epileptic spikes and electrical artifacts can produce extreme values. In a first step, the appropriate model order for GC-analysis was selected via Akaike Information Criterion using the LWR algorithm for faster computation. The full vector autoregressive (VAR) model was then estimated via ordinary least squares regression using the selected model order. By comparing the full VAR model to reduced VAR models that omit one of the predictors, this analysis yields an *F*-value for each directed comparison from the log-likelihood ratio:

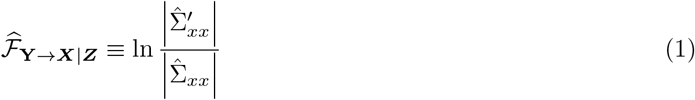

that can be read as *“the degree to which the past of Y helps predict X, over and above the degree to which X is already predicted by its own past and the past of Z”*^37^ 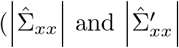 are determinants of the sample estimators of the residual covariance matrices for the full and the reduced model, respectively; for the univariate case these reduce to the variance). In other words: *F* denotes how much *Y* can add to the prediction of *X*. Importantly these values can be meaningfully compared^37^ and we here compare how much run 2 can add to the prediction of run 1, with how much run 1 can add to the prediction of run 2. This is interpreted as a measure of predictive recall (Fig. 2a).

#### Predictive recall channel selection

In order to separate channels that expressed neural learning of prediction from electrodes that didn’t express learning, a Gaussian Mixture Model with 2 components was fit onto the distribution of difference in *F* across channels. The underlying reasoning was that channels where learning was taking place should come from a distribution with a higher mean than channels where no learning was taking place (which should center around a zero mean). 31 channels were selected because the posterior probability of their observed difference in *F*-value was 10 times higher for the ‘effect distribution’ compared to the ‘null distribution’ (Supplementary Fig. 3b). Using the Yeo 17-network solution^41^, 15 of those electrodes were ascribed to network 4, 4 electrodes to network 14, 3 electrodes were ascribed to network 17, 2 to network 12 and 1 electrode to each of the networks 3, 6, 7, 8, 9, 13 and 16 (Supplementary Fig. 3c). Because the electrodes on which we observed predictive recall effects were located in cortical areas, we refer to them as cortical predictive recall (CPR) channels.

#### Connectivity analysis via Mutual Information

Connectivity analysis was performed via Gaussian Copula Mutual Information (GCMI). This method is rank-based, robust, and makes no assumptions about the marginal distributions of each variable, resulting in an estimate that is a lower bound of the true Mutual Information^57^. These properties make it a preferred method to account for potential outliers in the data due to residual epileptic spikes or inevitable artifacts because of the continuous nature of this data-set.

Specifically, conditional multivariate mutual information was used. In this, the mutual information between the high frequency amplitude (70 – 200 Hz) on all CPR-channels, on the one hand, and a given channel’s multivariate pattern (raw signal and amplitude in 6 frequency bands: delta < 4, theta 4 – 8, alpha 8 – 15, beta 15 – 30, low gamma 35 – 55 and high gamma 70 – 200) on the other hand, was computed at different lags: The multivariate channel pattern was shifted against the CPR-channels, from a 1.5 second lead to a 1.5 second lag. In order to take out spurious (and implausible) effects at lag zero^39, 40^, this lagged GCMI was conditioned on the multivariate channel pattern at zero-lag. For the connectivity analysis near event boundaries, data from 1-second windows around each potential boundary were considered and the whole procedure was repeated at different moments around the behaviorally recorded event boundary starting 3 seconds before the boundary and ending 1 second after the boundary. This shifting accounts for a potential mismatch between the moment of perception of an event boundary and the moment of button press (compare: Fig. 3a). Consequently this resulted in a 2 dimensional map that estimates shared information at different lags (first dimension) and at different time-points around the event boundary (second dimension) across all data within 1 second around these moments. For the connectivity analysis near peaks in predictive recall, data within 1-second windows around these peaks were considered, resulting in a 1 dimensional estimate of shared information at different lags.

#### Time-course of neural predictive recall

In order to derive a moment-by-moment time-course of predictive recall, we asked the question: Where does the model from run 2 predict run 1 better than the model from run 1 predicts run 2? In a first step, we derived the model prediction between runs (run 2 predicting run 1 and run 1 predicting run 2) on CPR-channels (see channel selection above). To this end, only the coefficients that describe the contribution of run 2 to the prediction of run 1 were multiplied with the data from run 2 and vice versa, i.e., if

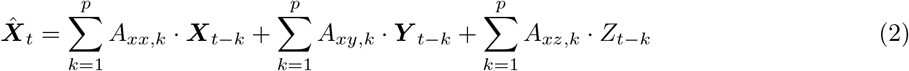

describes the full model predicting run 1 (*X*) at time-point *t* (where *Y* are the data from run 2, *Z* is the audio recording and *p* is the model order) a partial model-prediction from run 2 to run 1 was derived via

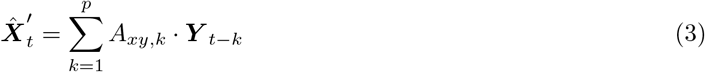

In order to assess when the model predicted the data accurately, the model-prediction was projected onto the data by taking the dot product between model-prediction and data across channels and then dividing by the number of channels. Specifically, we multiplied the model prediction 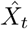 with the actual data *X_t_* at every channel and time-point (yielding positive values if the model prediction and the data were pointing in the same direction) and averaged across channels. The difference between the projected model that predicts run 1 from run 2 and the reverse model that predicts run 2 from run 1 was then smoothed with a moving average filter of 1 second width and was taken as a time-course of predictive recall in further analyses (Fig. 4a). To assess whether it would be useful to separately analyze the CPR-channels based on anatomical region (i.e., which Yeo network they belonged to), we also derived prediction accuracy separately for each channel (difference in model-times-data per channel) and correlated this measure between channels. Despite closer spatial proximity, pairwise correlations of time-courses at each channel (before averaging) were not significantly higher within anatomical networks than between networks (*t*(168) = 1.103, *p* = 0.272); we therefore decided to use the average time-course across all CPR-channels in further analyses, rather than grouping these electrodes by network.

#### Definition of peaks in time-courses

For peak detection, time-courses were first smoothed with a Gaussian window of 2 seconds width. Subsequently, the data were thresholded at the 95*^th^* percentile and grouped in clusters of neighboring points. Each cluster’s maximum was taken as a local peak. To relate the neural prediction time-course to event boundaries, peaks were taken from the second run of the story segmentation experiment and the neural time-course of prediction was locked to these peaks and averaged.

#### Correlation of behavioral predictive recall and neural predictive recall time-courses

Before correlating the behavioral data with the neural data, some behavioral participants were rejected from analysis. In the replication of the behavioral prediction experiment, participants were excluded if their vector of correct responses (coded as 0,1) had a cosine similarity <= 0.6 with the average prediction probability across all other *n* – 1 participants. This process was repeated until all participants were sufficiently similar, resulting in an exclusion of 9 participants in the naive condition and 7 participants that had heard the story before. To derive a time-course of behavioral prediction for correlation, a vector of the same length as the neural prediction time-course was created. This vector was holding the change in prediction probability of each word, at moments when a word was presented. The correlation between the neural predictive recall time-course and the behavioral predictive recall time-course was then computed only across moments where a word was presented (i.e. using a continuous probability vector ignoring silences between words). The latter was done to avoid confounding word-onset effects with differential effects between words (only the latter were of interest). The time axis was then shifted and the analysis was repeated to identify the lag that yielded the peak correlation.

### Statistical testing

#### Improvement in word prediction

In the behavioral word-prediction experiment and in its replication, the words’ probability of being predicted correctly was compared between the respective two groups (i.e., the group that had heard the story before, and the group that had not) with a dependent sample t-test, to test for an improvement in predictability across words. To test for improvement in word prediction performance across participants in the replication experiment, the probability of predicting the upcoming word correctly (ratio of correct predictions) was contrasted with an independent sample t-test between the 2 groups.

#### Story segmentation learning

Agreement was computed for each participant and each run by taking the cosine similarity of that participant’s response vector to the average response vector for the other participants. We then computed the average (across participants) of the difference in agreement across runs. This value was compared to the distribution of average differences under 1000 random permutations of run labels (Supplementary Fig. 1d). A simple binomial test was also used to assess the probability that agreement was improving across runs; this test evaluated the proportion of participants who showed improvement, under the null-hypothesis that the probability to improve was 0.5 for each participant. To assess significance of the lag between run 1 and run 2, the cross-correlation analysis was repeated 1000 times under random assignment of labels and the observed lag was compared to the random distribution of lags.

#### Neural learning

The presence of neural learning was tested conservatively by asking whether an overall effect was present in the data. To this end, the difference in *F*-values was averaged across all electrodes. Subsequently, the data were permuted 1000 times by applying a random sign-flip to each electrode’s *F*-value difference and then re-averaging across electrodes, resulting in a null distribution of averages. To ensure that the effect was not driven by a few patients, in another test, this permutation was done on a patient level (i.e., the same random sign-flip was applied to all of the electrodes from a given patient). Note that with 9 patients the maximal amount of distinct permutations is limited to 2^9^ = 512. In both tests, the true average difference across all electrodes was compared to the distribution of averages under random assignment of runs to determine significance. A *p*-value was computed as the number of differences that were larger under random permutation, divided by the number of permutations.

#### Phase shuffling to test cross-correlations

In order to statistically test the cross-correlations between the time-course of neural learning, and the time courses of behavioral learning, the neural time-course was phase shuffled 1000 times^58^ and the same cross-correlation was computed. A p-value was then computed for every shift of the time-axis as the number of correlations with phase-shuffled data that exceeded the true correlation at this lag, divided by the number of permutations. To correct for multiple comparisons, and thereby determine significance, a false discovery rate correction was applied across the time window of interest^42^. Whenever false discovery rate correction was applied, the largest significant p-value is reported as *p_FDR_*, hence all significant p-values are smaller than this threshold.

#### Mutual information near event boundaries

We computed mutual information (MI) between CPR-channels and hippocampal channels near event boundaries, and compared this to MI between CPR-channels and a visual region of interest (ROI), within a plausible range of lags and time-points around the event boundaries; this was done separately for run 1 and run 2. This statistical analysis assessed whether a cluster of high MI observed in the hippocampus was indeed larger than in the control ROI, where no such cluster was hypothesized. In a first step the hippocampal channels were compared to the control channels with a two-tailed independent sample t-test. 2-dimensional clusters were then formed by considering neighboring time-lag-points where t-values exceeded a cluster-forming threshold of a t-value that corresponds to an alpha threshold of 5% and the MI within these clusters was summed. The maximum cluster sum was then compared to the maximum cluster sums that were obtained after randomly swapping channels between hippocampus and control ROI 1000 times. A p-value was derived as the ratio of maximum clusters that exceeded the real maximum cluster under random permutations.

#### Peak locked analyses

In analyses that test for elevated values near peaks, statistics were assessed by repeating the same analysis 1000 times with randomly selected peaks; this includes the analysis of neural prediction locked to event boundaries and the analyses of mutual information locked to predictive recall peaks. In order to account for the auto-correlational structure of the neural data, the random neural prediction peaks were extracted from the phase-shuffled neural data. From the randomly generated peaks, a distribution was generated at every time-point around these random peaks and a p-value was derived as the proportion of random values that were higher than the observed value (i.e., based on the true peaks). The resulting p-values were corrected for multiple comparisons by controlling the false discovery rate^42^; the largest significant p-values are reported as *p_FDR_*.

#### Effect sizes

Cohen’s *d* effect sizes were computed for t-values from a dependent sample *t*-test as 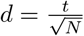 and for independent sample *t*-tests as 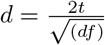

## Supplementary Information

1. MichelmannSI.pdf
  **Supplementary Text, Figures, Legends and Tables**
    Supplementary Text, Figures, Legends and Tables provide additional analyses, visual illustrations of statistics and electrode placement and information on patient details.
2. MichelmannSI.mp4
  **Supplementary Video**
    Video to Fig. 4a

### Supplementary Text

#### Supplementary Methods

##### Distribution of consensus on event boundaries between raters

In order to compare consensus between the first and second run of boundary norming, the distribution of agreement was compared between the 2 runs (agreement at each moment is the proportion of participants that marked that moment as an event boundary). In a first step, the quantiles of the distributions (across time) were plotted against each other in a Q-Q plot. We then analyzed the distribution of the data by binning the moments in the story into 232 bins, derived from the Freedman-Diaconis rule: 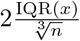. The difference in distribution was thereafter compared by assessing their kurtosis. Statistics on the kurtosis were computed on the difference between the first and the second run. The true difference was compared to the distribution of differences of 1000 randomly assigned labels; a p-value was derived as the ratio of differences that were bigger under such random permutation (Supplementary Fig. 1b, right).

##### Similarity in meaning of word predictions in the replication of the word-prediction experiment

In the main text, the behavioral analysis of prediction accuracy used a binary dependent measure (i.e., was the correct word predicted or not). Here, we explore a more continuous measure of accuracy, where we compute the cosine similarity between the Glove vector embeddings of the correct word and the predicted word. Specifically, we used the 300-dimensional 42B vectors that were pre-trained on common crawl^59^. These parametric estimates are arguably more sensitive than the binary measure used in the main text, insofar as they give partial credit if the predicted word is semantically similar to, but not the same as, the correct word (e.g., if the participant predicts ‘journalist’ instead of the correct word ‘reporter’). Statistically, the average cosine similarity between word embeddings of predicted and correct words was compared between the two groups (i.e., participants who had and had not heard the story before) with a dependent sample t-test. Additionally, each subject’s average cosine similarity between word embeddings of their predictions and the correct words was compared between the two groups with an independent sample t-test.

##### High frequency power around word onsets

To determine the peak response in high-frequency activity (70 – 200*Hz*) to word onset, we applied a multitapering approach with a window width of 200*ms* and three orthogonal Slepian tapers, i.e., a spectral smoothing of 10*Hz*. Power was estimated in steps of 5*Hz*, omitting the 120*Hz* and 180*Hz* band (as per harmonics of line noise). Averaged power was subsequently locked to the onset of words, data were z-scored across time and averaged across all words and channels that displayed a prediction effect.

##### Additional correlations between behavior and neural predictive recall

As an additional way of relating event boundaries to neural predictive recall, we computed the cross-correlation of these two measures. For this analysis, 13 behavioral subjects who provided event boundary ratings were rejected as being outliers (based on a cosine similarity <= 0.15 to the average response vector across all other subjects on either of the runs). The time-course of agreement (of button-presses indicating event boundaries) was down-sampled to the neural time-course of prediction and correlated. The neural time-course of prediction was shifted against the behavioral time-course to identify the lag that yielded the peak correlation. In a separate analysis, we also examined the correlation between neural predictive recall and the change in behavioral next-word prediction accuracy, this time using the cosine-similarity behavioral prediction measure described above (instead of the binary measure used in Fig. 4c in the main text). Cross-correlations were compared statistically to 1000 cross-correlations with phase-shuffled data; p-values were derived as the proportion of random correlations that were higher than the true correlation; p-values were corrected for multiple comparisons by controlling the false discovery rate^42^.

#### Further evidence for increased consensus on event boundaries upon one-shot learning

We predicted that subjects would have a better understanding of the underlying event structure of the story on the second run, and therefore would agree more about event boundaries on the second compared to first run, despite not knowing of each other’s responses and not receiving feedback about the accuracy of their responses. This change in the distribution of agreement is visible in a quantile-quantile plot (Supplementary Fig. 1a): The deviation from a diagonal line reflects a relative over-representation of moments of high agreement on the second run. Statistically, more accurate responses on the second run also result in more observations in the tails of the agreement distribution. In keeping with this, kurtosis on the second run (24.482) was higher than the kurtosis on the first run (17.94, Supplementary Fig. 1b, left) and this difference (6.512) was significantly higher than expected by chance *p* = 0.036, i.e., under random assignment of run-labels (1000 permutations, Supplementary Fig. 1b, right).

#### Further evidence for one-shot learning of word predictions

In the replication experiment of behavioral prediction learning, we could analyze the quality of individual word predictions by computing the similarity between predicted word and correct word via the cosine-similarity between their 300 dimensional Glove vector embeddings (pre-trained on common crawl)^59^. This average similarity was higher in the group that had listened to the story before (*t*(959) = 34.967, *p*< 0.001, *d* = 1.127, Supplementary Fig. 1d, purple line), providing converging evidence that participants were learning to predict the next word based on prior exposure to the story. In the replication experiment we could also compare subjects’ prediction performance between groups. Subjects’ vector embeddings of their predictions were more similar to the correct words (*t*(98) = 6.512, *p*< 0.001, *d* = 1.316) if they had heard the story before.

#### Cross-correlation between neural predictive recall and event boundary time-course

To further test whether predictive recall encompasses the structure of the story, thereby allowing for the anticipation of boundaries, we assessed cross-correlation between the behavioral time-course of agreement (Fig. 1a) and the neural time-course of memory-based prediction (Fig. 4a). When we correlated the average time-course of agreement from the first and second behavioral run with the neural time-course of memory-based prediction, we obtained a maximum correlation of *r* = 0.217 at −1.69 seconds (i.e., before boundary detection); note that this negative lag is expected because a behavioral response can only happen after an event has been neurally registered. This correlation was significantly higher than what we obtained with phase-shuffled time-courses of memory-based prediction; several lags survived a false discovery rate multiple comparison correction (*p_FDR_* = 0.002, 1000 permutations, Supplementary Fig. 4a). If behavioral learning leads to a more accurate representation of an underlying event structure, we further expected that the time-course of agreement derived from the second behavioral run should correlate slightly better with the neural data. Indeed, with the second behavioral run (*r* = 0.221, peak: −1.61*s*) the maximum correlation was slightly higher than the with the first behavioral run (*r* = 0.197, peak: −1.76*s*), however, only a statistical trend was observed when this difference (0.024) was compared to differences of randomly re-sampled behavioral groups (*p* = 0.063).

#### Further analyses linking neural predictive recall to one-shot learning of story content

In the main text, we report that the maximum correlation between neural predictive recall and (behavioral) predictive recall of individual words in the “replication” experiment (*r* = 0.113) was found at 590*ms* after word onset (*p_FDR_* = 0.006, Fig. 4c, fuchsia). When we repeated this analysis using cosine-similarity to measure predictive recall of individual words (instead of the binary measure used in the main text), the maximum correlation (*r* = 0.127) was found at 320ms after word onset *p_FDR_* = 0.007 (Supplementary Fig. 4b, purple). This more sensitive analysis therefore converges with the lag found in the original behavioral prediction experiment (Supplementary Fig. 4b, turquoise).

#### Gamma response to word onset on CPR-channels

The neural high frequency response (70 – 200Hz) to word-onset peaked at 320ms (run 1) and at 389*ms* (run2, Supplementary Fig. 4b). These lags correspond closely to the lag obtained in the preceding analysis (i.e., when we looked for the maximal cross-correlation between the time courses of neural and behavioral predictive recall), suggesting that neural learning entailed predictions about individual words via anticipation of the neural high frequency response to word-onset.

#### Feature analyses of Multivariate Mutual Information related to predictive recall

In order to clarify which frequency bands were most relevant for the increased multivariate mutual information at moments of predictive recall (compare Fig. 5a, right), we repeated this analysis omitting each feature in the hippocampus (Supplementary Fig. 5a). This analysis demonstrates how much the overall effect diminishes if each feature is removed from the analysis. We further recomputed the MI for each hippocampal feature in isolation (Supplementary Fig. 5b). Mostly the raw signal and low frequencies (< 35*Hz*) shared information with cortex.

### Supplementary Figures

**Supplementary Figure 1.**
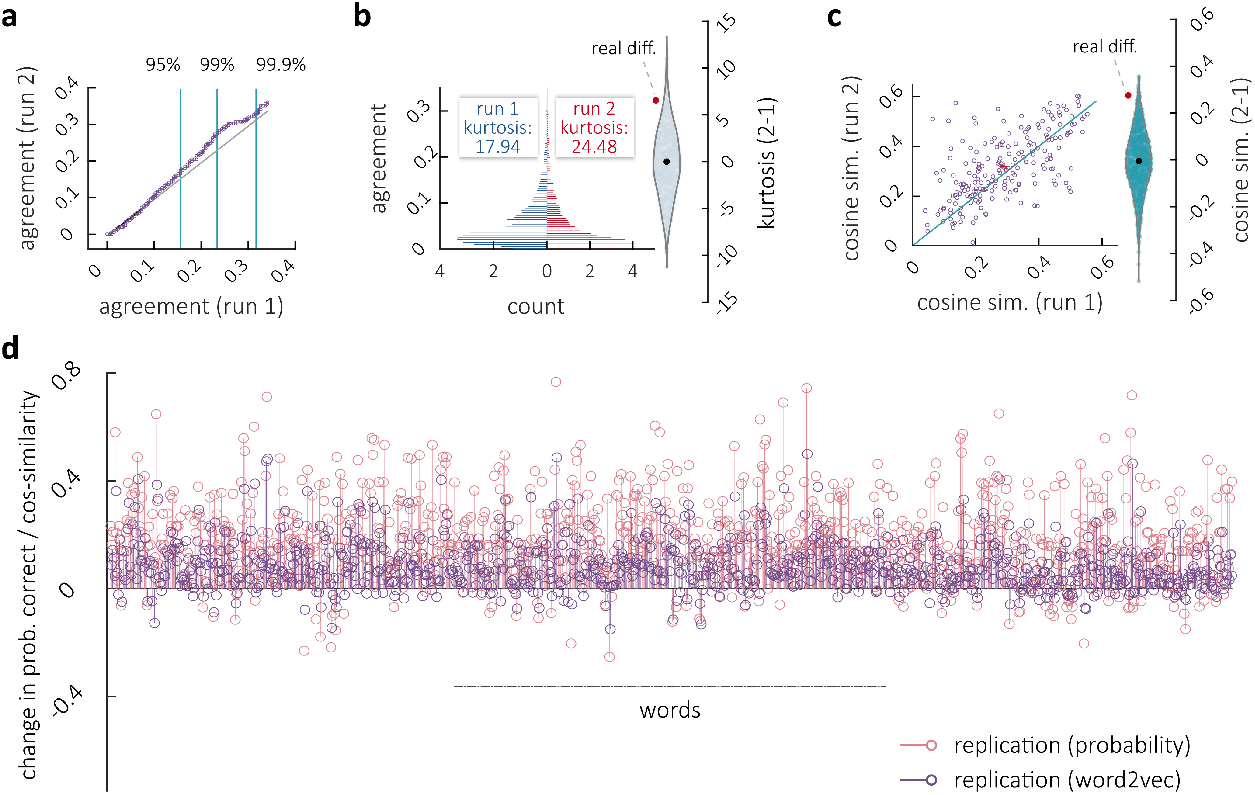
Behavioral analyses demonstrating predictive recall. **a.** Quantile-quantile plot comparing the distribution (across time) of agreement on event boundaries (compare Fig. 1a) between runs. The deviation from a diagonal line indicates a relative over-representation of moments of high agreement on the second run. **b.** Histograms depicting the distribution (across time) of agreement on the first run (left, blue) and on the second run (right, red). Higher kurtosis on the second run indicates that more values fall in the tails of the distribution. The violin plot displays the run-differences in kurtosis under random label permutation (gray) and the true difference (red dot) **c.** Similarity of each subject’s response vector to the agreement across all others (same as Fig. 1b). Purple dots indicate individuals: subjects above the diagonal increase in similarity to others on the second run. The average increase (red cross) indicates consensus learning. The violin plot displays the average run-difference in cosine similarity to others under random label permutation (turquoise) and the true difference (red dot) **d.** Performance difference between groups that had listened to the story and naive participants in prediction-measures for upcoming words in the story. Difference in prediction-probability (fuchsia replication) and cosine-similarity between predicted and correct word (purple experiment 2) are higher after a single exposure.

**Supplementary Figure 2.**
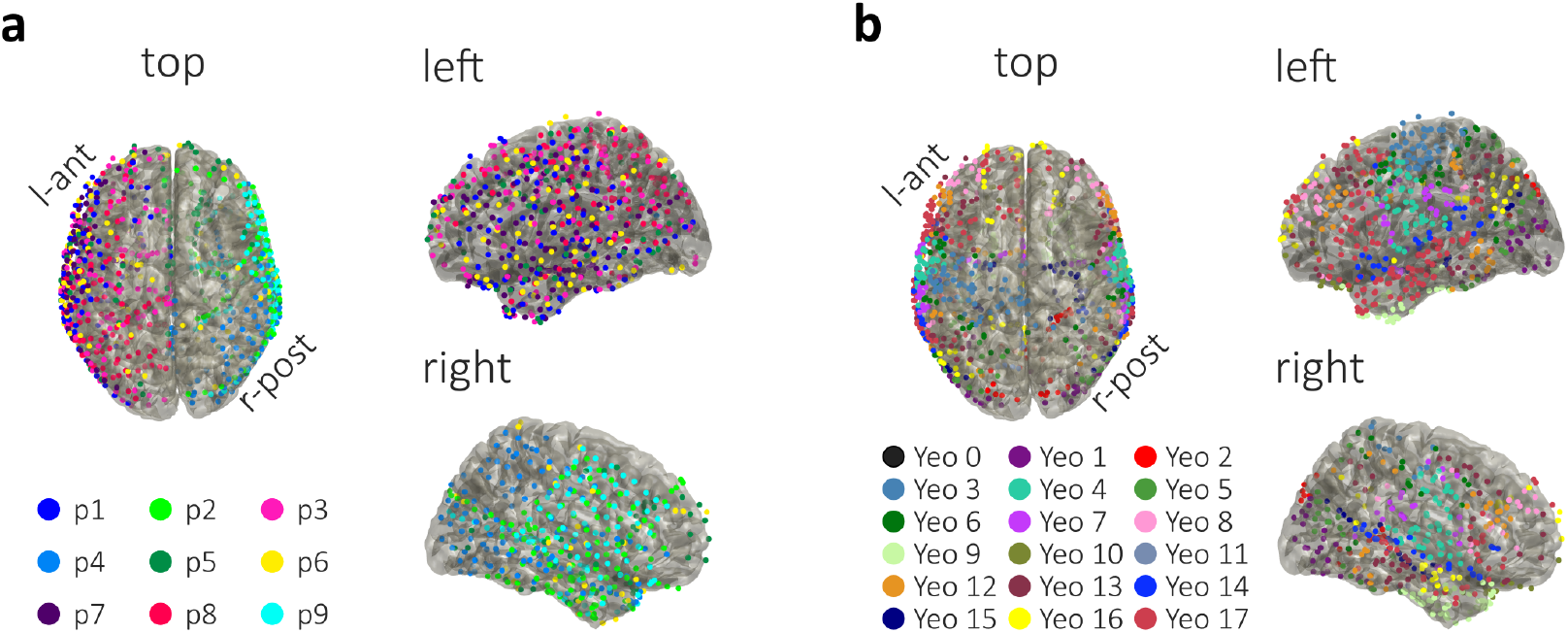
Electrode placement and anatomical labels. **a.** Electrode placement across nine patients showing extensive coverage across both hemispheres. **b.** Anatomical labels of electrodes according to the yeo 17 network solution (see: Online Methods).

**Supplementary Figure 3.**
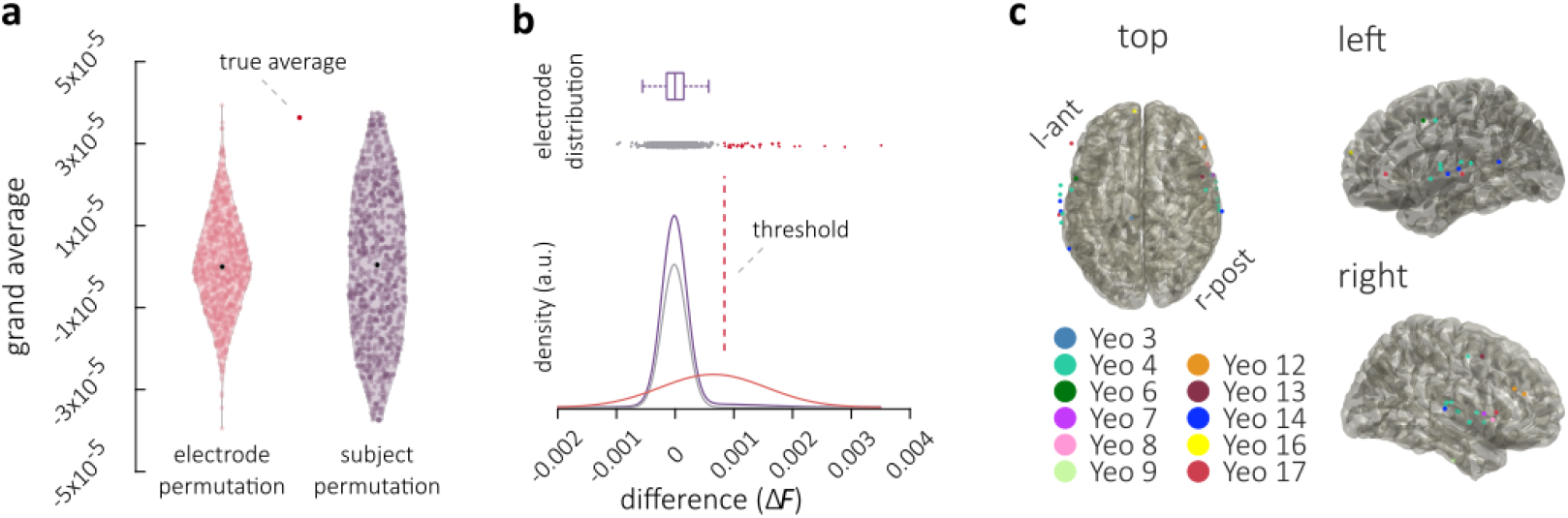
Statistics of neural predictive recall, electrode selection and labeling for ‘CPR-channels’. **a.** Differences in *F*-values measuring predictive recall (from the Granger Causality analysis) were averaged across all channels (true average, red dot). The pink violin plot displays these average values under random assignment of run-labels per channel. The purple violin plot displays average values under random assignment per subject (compare Fig. 2). **b.** Gaussian-Mixture-Modeling of the distribution of differences with 2 Gaussian distributions. Electrodes were selected as displaying predictive recall if they were 10 times more likely to belong to the distribution with the higher mean. The mixed distribution is purple, the gray Gaussian represents the null-distribution and the red Gaussian the effect-distribution (scaled to an AUC of 1 for visibility). Dots are electrodes in the color of their assigned distribution. **c.** Effect-electrodes are colored according to their position corresponding to the 17 network parcellation of the Yeo-atlas.

**Supplementary Figure 4.**
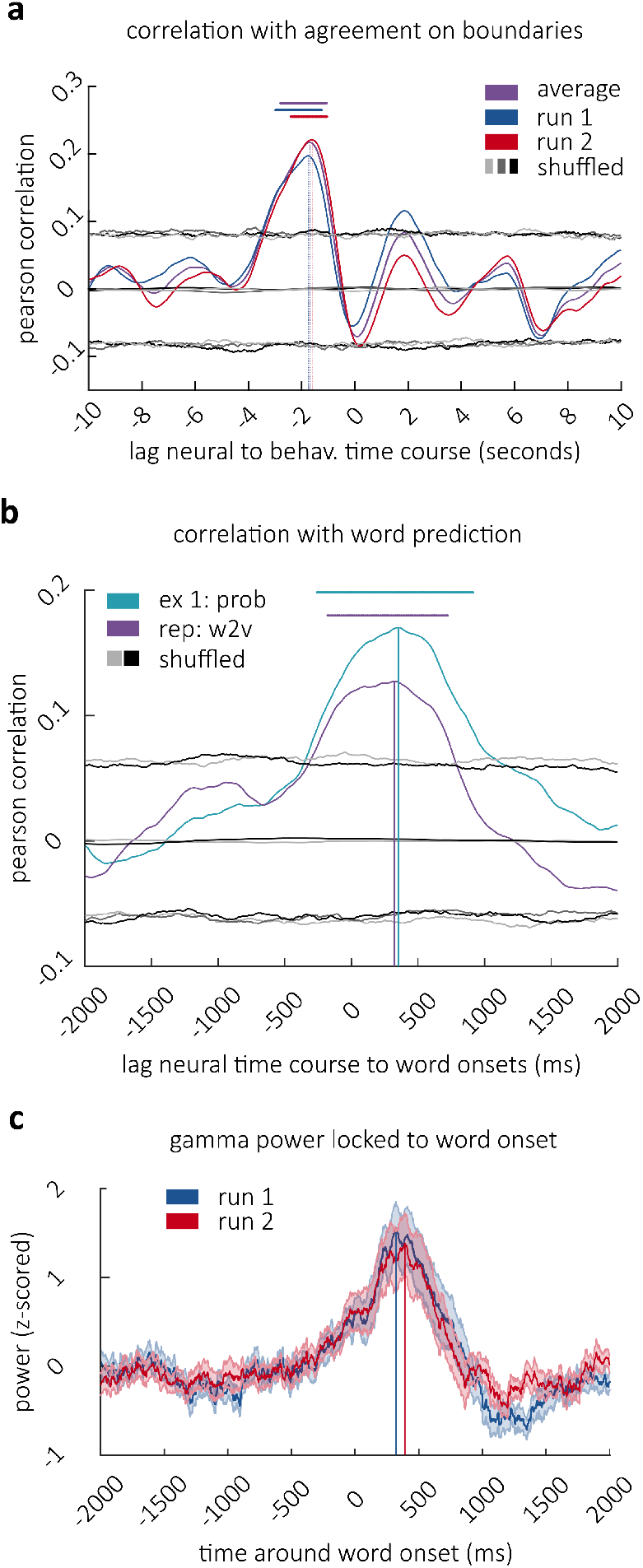
Cross-correlation between neural prediction and event boundary time-course and gamma response to word onset on CPR-channels. **a.** Cross-correlograms of the neural time-course of predictive recall with the time-course of agreement on boundaries (compare Fig. 1a, with average between behavioral runs: purple, with behavioral run 1: blue, with run 2: red), vertical lines mark maxima. Gray-scaled lines are 5*^th^* and 95*^th^* percentile of cross-correlations with phase-shuffled neural data. Horizontal lines mark significance (fdr-corrected). **b.** Correlation between the behavioral measures of word-prediction learning (compare Fig. 1e, turquoise, Supplementary Fig. 1e, purple) and the neural time-course of predictive recall at different time-lags. Line in turquoise shows correlation between the probability of correct prediction in experiment 1, the purple line represents the correlation with the difference in cosine-similarity from the replication, which is arguably a more sensitive measure of prediction learning. Gray lines are 5^*th*^ and 95^*th*^ percentile of correlations under random assignment of performance to words for the respective measures. Horizontal lines mark significance, vertical lines mark peaks. **c.** Average response (+ − *SEM*) in the gamma band to word onset in run 1 (blue line) and run 2 (red line) on CPR-channels. Vertical lines mark peaks.

**Supplementary Figure 5.**
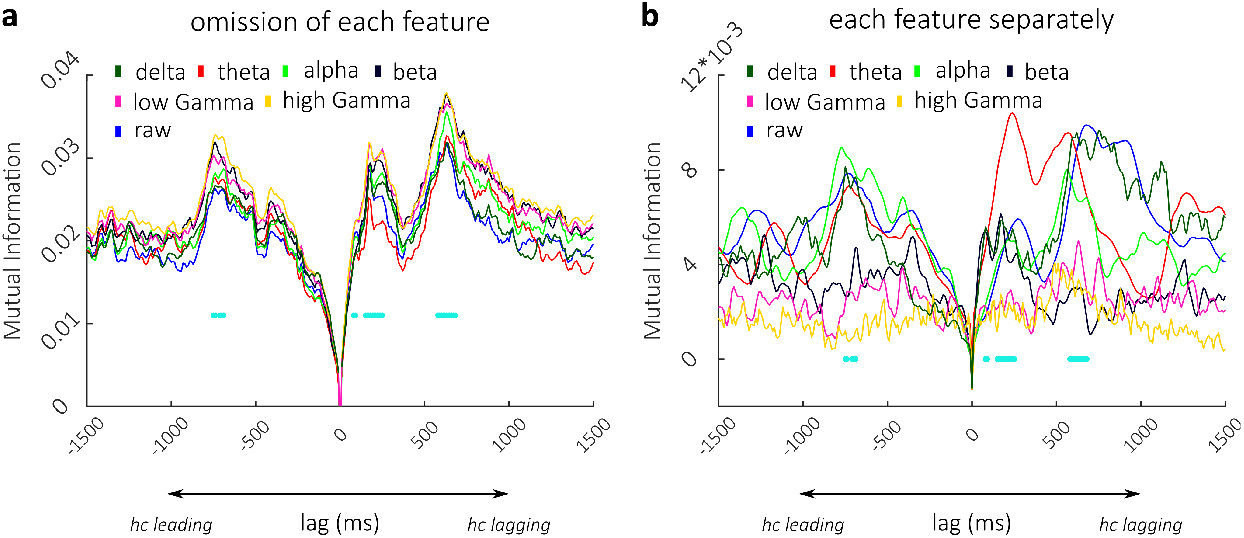
Feature by feature re-analysis of hippocampo-cortical connectivity at moments of predictive recall on run 2. **a.** Conditional Multivariate Mutual Information at different lead/lag to CPR-channels at neural predictive recall events was re-analyzed on run 2 omitting each hippocampal feature once. These data show the contribution of features to MI via reduction in MI. **b.** Conditional Multivariate Mutual Information at different lead/lag to CPR-channels at neural prediction events was re-analyzed on run 2 using each hippocampal feature on its own. These data show the contribution of features to MI, specifically, how much MI is obtained by using each feature on its own. Horizontal lines depict points of significance from Fig. 5a, right in turquoise.

### Supplementary Tables

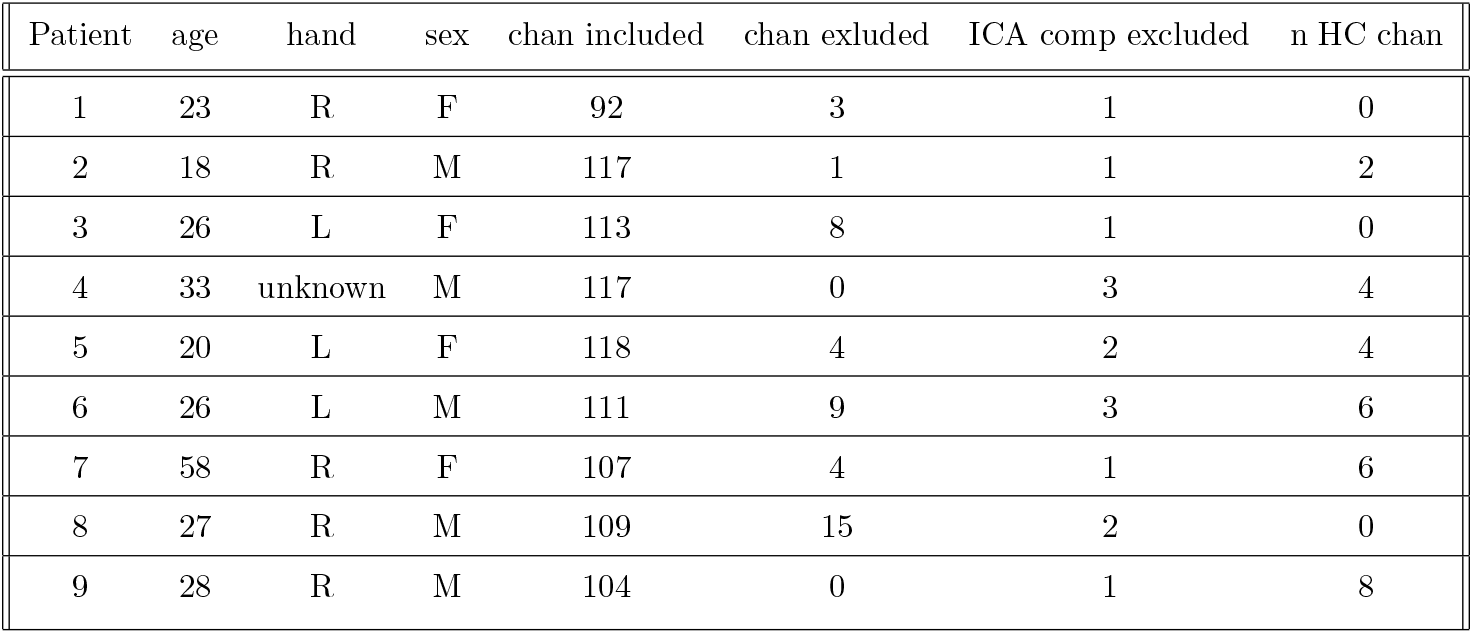
Details on recorded patients

